# Focal Neurostimulation of Calcium Signaling and Dopamine Release in Human Dopaminergic Neurons Using Megahertz-range Single-Pulse Focused Ultrasound

**DOI:** 10.64898/2026.02.14.705923

**Authors:** Sarvenaz Khodayari, Ivan Suarez-Castellanos, Magali Perier, Tom Aubier, Parastoo Hashemi, Alexandre Carpentier, Stephane Marinesco, W. Apoutou N’djin

## Abstract

Focused Ultrasound (FUS) neurostimulation has increasingly attracted attention given its ability for localized targeting and non- to minimally-invasive capacity. However, the understanding of the biological and neurochemical mechanisms triggered by this neurostimulation modality remains limited. Indeed, further progress of this technology could benefit from spatiotemporal evaluation of neurotransmitter secretory activity resulting from FUS stimulation. Recently, we demonstrated *in-vitro* the ability of FUS to evoke calcium (*Ca*^2+^) waves, in a mixture of human neuronal and glial cells (standard cells), with single-pulse megahertz-range FUS compatible with a transdural approach relying on an intracranial FUS implant. In the present work, we investigated the ability to evoke *Ca*^2+^ signaling and dopamine (DA) release by single-pulse megahertz-range FUS stimulation of cultured human dopaminergic neurons *in-vitro*. A hybrid-platform integrating a custom-made FUS implant prototype (concave spherical transducer, Ø and focal distance: 15 mm) with real-time *Ca*^2+^ fluorescence microscopy imaging (FMI), and fast scan cyclic voltammetry (FSCV) was constructed to evaluate FUS-evoked *Ca*^2+^ signaling and concurrent DA release. *Ca*^2+^ and DA releases evoked by FUS on dopaminergic neurons (DA neurons) were compared to those evoked in standard cells. Application of single-pulse FUS (frequency: 5.11 MHz, pulse duration: 700 µs, spatial average pulse average intensity, *I*_*sapa*_: 19.59 ± 4.12 W.cm^−2^) was shown to causally mobilize *Ca*^2+^ dynamics in both cell types. Immediate (< 1 s) responses were focally evoked in cell clusters of 290 µm in diameters; corresponding to the −3 dB FUS focal diameter. However, while standard cells exhibited continuous and omnidirectional delayed propagating dynamics following FUS stimulation, DA neurons showed more spatially sparse responses. The FUS-induced *Ca*^2+^ activity in DA neurons was accompanied by DA release detected by FSCV, but not in standard cells. This study demonstrates that single-pulse megahertz-range FUS induced with an intracranial implant prototype smaller than conventionally used FUS devices (transcranial applications) can evoke *in-vitro* intracellular *Ca*^2+^ activity while stimulating DA release from dopaminergic neurons, underscoring its potential as a neuro-stimulation/-modulation tool for targeting dopaminergic circuits in various pathologies.

## Introduction

Neurological and neuro-degenerative disorders represent a growing global health challenge, and their incidence is predicted to rise significantly as the world’s population ages^1^. These conditions gradually impair neuronal structure and function, leading to chronic and progressive deficits in motor, cognitive, and behavioral capacities^2^. Neurological and psychiatric diseases sometimes result from degenerating or hypoactive groups of neurons within the central nervous system (CNS); in these cases, stimulating these neurons can improve the patients’ state. For example, electrical stimulation of the subthalamic nucleus results in significant improvement of motor symptoms in Parkinson’s disease (PD) patients^3^. Deep brain stimulation (DBS) can also be effective in treatment resistant depression^4^. Noninvasive neuro-modulation/stimulation techniques are in high demand for improving these therapeutic strategies without the risks associated to brain surgery^5,6^. These interventions include transcranial alternatives such as transcranial magnetic stimulation (TMS) or direct and alternating current stimulation (tDCS and tACS). By modulating or stimulating specific brain regions or neural pathways, neuromodulation can provide symptom relief, enhance neuroplasticity, and, in some cases, reduce the need for pharmacological treatment.

Focused ultrasound (FUS) neuromodulation has emerged as a promising noninvasive or minimally invasive technique for the targeted modulation of neural activity^7,8^. By adjusting its acoustic parameters, FUS can elicit precisely controlled thermal or mechanical effects within specific anatomical regions, allowing for high spatial selectivity^8,9,10^. These bioeffects have been increasingly harnessed to develop therapeutic strategies for a variety of pathological conditions, including those involving the CNS^7,8^. In the context of neurological disorders, numerous studies have investigated the potential of FUS to modulate or stimulate neural circuits in a spatially and temporally precise manner. Following the definitions proposed by Gavrilov’s foundational work^11^, specific terminologies can be used to clarify the outcomes of the various modalities. While neuromodulation refers to the process of influencing the inherent activity of a neural structure by modifying its state – oftentimes using low energy but prolonged exposures to the stimuli - neurostimulation describes the initiation of neural activity consequent to a short and rarely repeated stimulus^11^. Nevertheless, despite substantial progress in the field, the fundamental mechanisms underlying FUS-induced neural activation and regulation remain incompletely understood^7,10^, particularly regarding its cell-type–specific and circuit-level effects across different neural populations. Elucidating how ultrasound influences distinct neuronal networks is therefore essential for advancing its translational potential in both experimental and clinical neuromodulation. Among these networks, the dopaminergic system represents a particularly compelling target due to its central role in motor control, reward processing, and neuropsychiatric disorders^12,13^. Dopamine (DA) is a monoamine neurotransmitter derived from the amino acid tyrosine; it plays an important role in several essential functions in the CNS, including motor control, learning and attention, motivation and reward. DA is mostly produced by neurons in the substantia nigra and ventral tegmental area. DA is released through calcium (*Ca*^2+^)-dependent vesicular exocytosis^14^.

Abnormal release of DA can result in a variety of neurological diseases such as PD, Schizophrenia and depression. Neuromodulation strategies, including FUS, influence the release of DA and other neurotransmitters (e.g., GABA, glutamate), which may help to restore a more physiological chemical environment. Recent studies demonstrate that transcranial FUS can non-invasively modulate neurotransmission, for instance by reducing extracellular GABA levels in rats without affecting glutamate. Similarly, FUS has been shown to increase DA and serotonin release in the frontal cortex, and in humans, theta-burst ultrasound stimulation (involving repetitive-pulse FUS sequences) selectively reduced GABA concentrations and enhanced functional connectivity, supporting its potential as a neuromodulator tool. Previous studies have also evaluated and demonstrated the capabilities of FUS neuromodulation/ neurostimulation strategies to effectively evoke the release of DA from dopaminergic cells or circuits.

Tian Xu et al.^15^ demonstrated that 1 MHz transcranial ultrasound stimulation can modulate neural activity and improve motor symptoms in PD, highlighting its potential as a non-invasive neuromodulator therapy. However, none of these studies have sought to investigate and describe the spatiotemporal characteristics of DA secretory responses to FUS neurostimulation sequences, and a variety of repetitive-pulse FUS protocols have been proposed. Characterizing the secretory dynamics of DA induced by elementary FUS protocols involving single FUS pulses may serve as building blocks for the design and development of more complex, repetitive-pulse FUS-based therapeutic strategies targeting the underlying causes of abnormal secretory activity in diseases such as PD. Our team has explored the use of such single-pulse FUS sequences, in the context of a transdural approach relying on an intracranial FUS implant (Fig. 1(a)), to stimulate and analyze causal, targeted neural responses across models of varying anatomical complexity. Recent work by Aubier et al.^16^ demonstrated that single-pulse megahertz-range FUS can induce *Ca*^2+^ fluxes and concurrent glutamate release from glial cells, whereas Suarez-Castellanos et al.^17^ showed that the same stimulation paradigm can evoke excitatory postsynaptic potentials (EPSPs) in *ex-vivo* hippocampal circuits. However, these studies did not characterize the spatiotemporal dynamics of FUS-induced neurotransmitter release from targeted neurons, nor did they distinguish between direct glial and neuronal activation mechanisms.

**Figure 1.**
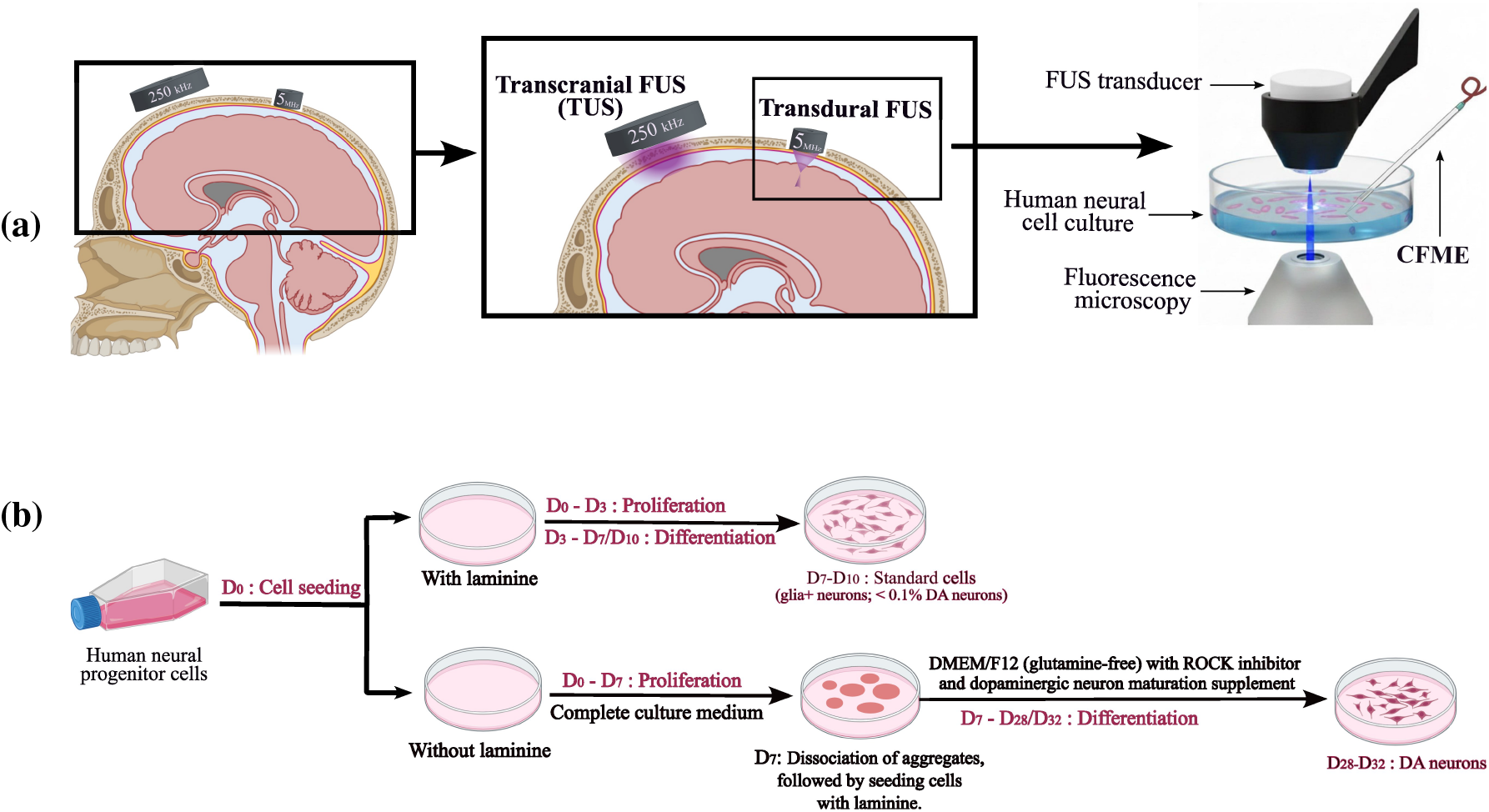
Single-pulse focused ultrasound (FUS) neurostimulation studies on an *in-vitro* model of human DA neurons using a transdural intracranial FUS technique. (a) From left to right an illustration of a FUS stimulation strategy using a transdural FUS cranial implant to provide very targeted and persistent stimulation schemes. Against the transcranial FUS (TUS), which shows a wide targeted area without special spatial precision. *in-vitro* model of employing FUS prototype and simultaneous monitoring using carbon fiber microelectrode (CFME). (b) *In-vitro* cell preparation stages: standard cells vs. dopaminergic (DA) neurons.

Several neuromonitoring techniques can be used to track the release of DA neurotransmitters resulting from FUS neurostimulation. Among such methods, enzyme assays^18^, mass spectroscopy and liquid chromatography are popular approaches for evaluation of monoamine content from neurochemical mixtures^19^. However, despite their powerful quantitative detection, these techniques remain limited by low temporal resolution, and difficulty of applying them *in-situ*. On the other hand, electrochemical techniques offer high temporal resolution, *in-situ* detection, and excellent sensitivity for monitoring dynamic DA release, in addition to lower costs^20^. Consequently, such techniques would be well suited to monitor FUS-induced neural activity in the form of DA secretion. Characterizing the secretory dynamics in time and space is essential, not only for elucidating the fundamental mechanisms of FUS-induced neuromodulation, but also for optimizing stimulation parameters to achieve targeted and predictable modulation of dopaminergic activity.

Fast-scan cyclic voltammetry (FSCV) is an electrochemical detection method that combines with carbon-fiber microelectrode (CFME) and provides a sub-second resolution to detect the activity of neurotransmitters like DA and serotonin^21,22^. The combination of FUS neurostimulation and FSCV neuromonitoring strategies offers a powerful experimental framework for achieving targeted stimulation and real-time monitoring of neural activity with high spatial and temporal resolution. This combined approach holds significant promise for elucidating the mechanisms underlying neurostimulation and for advancing the development of precise, safe neuromodulatory therapies for neurodegenerative and neuropsychiatric disorders.

In this *in-vitro* study, we investigated the feasibility of evoking *Ca*^2+^ signaling and DA release in DA human neuron cultures using single-pulse megahertz-range FUS. A novel custom-made FUS implant prototype was specifically developed which included an operative channel to integrate with CFMEs and FSCV. This hybrid setup, combined with fluorescence microscopy imaging (FMI), enabled simultaneous real-time monitoring and post-analyses of the spatiotemporal dynamics of intracellular *Ca*^2+^ signaling and extracellular DA release evoked by FUS neurostimulation. The specific characteristics of DA neuron responses to single-pulse FUS were studied in comparison with those produced in a standard differentiated neural culture containing both glial and neuronal populations involving < 0.1% of DA neurons.

## Materials and methodes

### a. Cell cultures preparation: Standard cells versus Dopaminergic neurons

All experiments were performed on standard and DA cultures obtained by differentiating the ReN-Cell ® VM cell line (SCC008, Merck, UK), a human neural progenitor cell line with the ability to differentiate into neurons, astrocytes and oligodendrocytes (Fig. 1(b)). Cells were seeded at a density of 30,000 cells per cm^2^ in 35 mm-diameter Petri dishes (Cellview, Greiner bio-one, Austria) coated with 2 mm-thick PDMS (Polydimethylsiloxane, Sylgard 184, Dow, USA). The PDMS was coated with 20 µg.ml^−1^ laminin (L2020, Sigma-Aldrich, Germany) to enhance cell adhesion.

In **Standard differentiated cultures**, cells were cultured with ReNcell ® NSC Maintenance Medium (SCM005, Merck Millipore, Germany), to which epidermal growth factor (EGF), (GF001, Merck, UK) and fibroblast growth factor (FGF), (GF003, Merck, UK) both at 20 ng.ml^−1^, were added. After 3 days, growth factors were removed to initiate differentiation. This medium did not allow for cellular differentiation of progenitors and was referred to as “complete” culture medium^23^. The obtained standard differentiated neural culture contained both glial and neuronal populations involving <0.1% of DA neurons. Cells were used between *D*_7_ and *D*_10_. This standard culture was recently used by our group to prove the ability of single pulse megahertz-range FUS stimulation in evoking *Ca*^2+^ signaling^16^. In the present study, the standard culture was used as a reference for comparison with dopaminergic neuron cultures.

For **dopaminergic differentiation**, cells were first seeded in an uncoated container at 30,000 cells per cm^−2^ in complete medium, allowing them to form aggregates during the initial 7 days. The aggregates were then partially dissociated using Accutase (A1110501, Gibco, USA) and reseeded onto laminin-coated, 2 mm-thick PDMS substrates to ensure proper adhesion. From this stage onward, cells were maintained in glutamine-free DMEM-F12 medium supplemented with 5 µM ROCK inhibitor (SCM075, Sigma-Aldrich, Germany) and 1× Dopaminergic Neuron Maturation Supplement (A3147401, Gibco, USA). The culture medium was renewed every 3 days. Dopaminergic neuron cultures were used between *D*_27_ and *D*_30_.

### b. Characterization of Standard and Dopaminergic cultures by immunological labeling

Immunocytochemistry was performed on cells seeded at 30,000 cells per cm^−2^ onto 13 mm Nunc Thermanox coverslips (10252961, Fisher Scientific, USA). Before staining, the cells were rinsed twice for 3 min with 500 µL of Phosphate-Buffered Saline (PBS 1X, Fisher Scientific, USA), followed by fixation with 500 µL of 4% paraformaldehyde (15670799; Fisher Scientific, USA) in PBS 1X for 30 min at room temperature; this step was followed by two additional rinses with PBS 1X. Permeabilization was performed using 0.2% Triton X-100 (11488696, Fisher Scientific, USA) in PBS for 10 min at room temperature, and cells were again rinsed twice with PBS 1X. Non-specific binding was blocked in PBS for 30 min at room temperature with 500 µL of 1% bovine serum albumin (BP9702-100; Fisher Scientific, USA).

Primary antibodies were diluted in PBS containing 1% BSA and incubated overnight at 4 °C, followed by 1 h at room temperature. The following primary antibodies were used: mouse anti-Nestin (MA1110, Invitrogen, USA; 1:1000), rabbit anti-tyrosine hydroxylase (TH; AB152, Merck, UK; 1:1000), and rabbit anti-glial fibrillary acidic protein (GFAP; AB5804, Merck, UK; 1:3000). After two rinses with PBS 1×, cells were incubated for 1 h at room temperature with the appropriate fluorophore-conjugated secondary antibodies (diluted 1:1000 in PBS containing 1% BSA): species-specific anti-mouse Alexa Fluor 546 (10082322, Fisher Scientific, USA) for Nestin, and anti-rabbit Alexa Fluor 488 (10729174, Fisher Scientific, USA) for TH and GFAP.

Hoechst 33258 (5 µM final concentration; ab228550, Abcam, UK) was used as a colorant for the nuclei, prepared as a 1:4000 dilution in PBS containing 1% BSA, and incubated for 15 min at room temperature. This step was followed by two final washes in PBS 1X, coverslips were mounted using Fluoromount mounting medium (F4680, Merck, UK). Immunostained samples were imaged with an inverted fluorescence microscope (Eclipse Ti-S, Nikon Corp., Japan), with a ×20 magnification equipped with an ORCA-Fusion CMOS Camera (C14440-20UP, Hamamatsu Photonics, Japan). They were then colorized and merged with ImageJ free access software (National Institutes of Health, Bethesda, MD).

### c. *In-vitro* neurostimulation platform : novel FUS prototype coupled to FSCV

A novel custom-made FUS implant prototype was specifically developed with an operative channel to integrate with CFMEs and FSCV neuromonitoring in an *in-vitro* set-up, while remaining compatible in size (Ø= 15 mm, height = 15 mm) with future intracranial transdural FUS neurostimulation approaches (Fig. 2(a)). The spherical ceramic FUS transducer (C-21, Ø= 15 mm, radius of curvature = 15mm; Fuji Ceramics Corp., Japan), providing an active surface of 1.76 cm^2^, perforated at its center (hole Ø= 2 mm), was sealed in a SLS 3D-printed nylon packaging (PA12, Xometry Europe, Germany) to form an operative channel allowing the passage of a CFME biosensor (Fig. 2(d)). A function generator (Tektronix AFG1062, Tektronix Inc. USA) and a 55 dB power amplifier (1140LA, Electronics & Innovation (E&I), USA) were employed to drive the FUS prototype. The experimental platform was built on an inverted microscope (Eclipse Ti-S, Nikon Corp., Japan) placed inside a Faraday cage sitting atop an anti-vibration table (CleanBench Gimbal Piston, TMC, USA). The setup was also equipped with a set of 2 automatic micro-manipulators (MPC-200, ROE-200, MP-225, Sutter Instruments, USA) for precise placement of the FUS prototype and the CFMEs. Customized holders were 3D printed to attach the FUS prototype to the micromanipulators while holding its active surface facing downwards to generate FUS vertically towards the sample holder and objective of the microscope.

**Figure 2.**
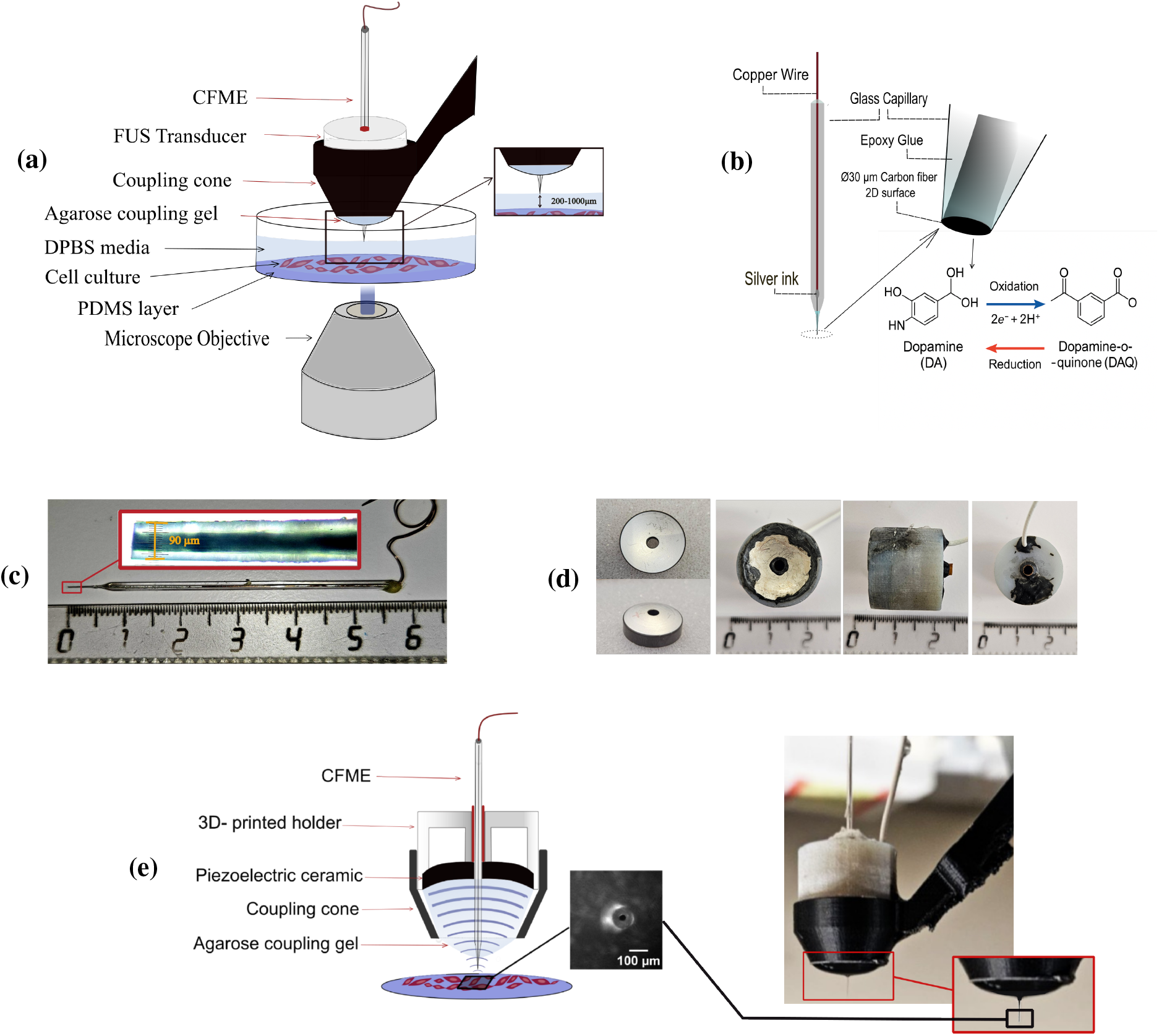
FUS stimulation and FSCV setups, shown separately and integrated together. (a) Schematic of the experimental setup illustrating the integration of the FUS transducer with the CFME. (b) Schematic of the CFME and its active 2D surface, including the DA redox reaction under applied FSCV, showing the oxidation of DA to DA-o-quinone (DAQ) and its subsequent reduction back to DA. (c) Optical image of the CFME. (d) From left to right: perforated spherical piezoelectric element (15 mm diameter, 15 mm curvature radius); final assembled transducer; and the piezoelectric element mounted inside the 3D-printed holder. (e) Left to right: Schematic crosssectional view of the transducer positioned within the 3D-printed coupling cone, and microscope image of the CFME 2D tip placed on top of cultured cells; side view of the transducer mounted in the 3D-printed coupling cone, filled with agarose gel for acoustic coupling.

For the *in-vitro* experiments, to maintain acoustic continuity between the FUS transducer and the neural cell monolayer in the cell medium-filled Petri dish, a 0.4% Agarose coupling gel (A9539 Sigma-Aldrich, USA) was cast in a 3D-printed SLS nylon coupling cone (PA12, Xometry Europe, Germany) prior to the experiments. The coupling cone was integrated in 3D printed piece connecting the FUS prototype to the micromanipulators. In addition, all experiments were performed in DPBS containing calcium (14040091, Gibco, USA), which provides a pH-neutral medium compatible with FSCV recordings. For simultaneous FUS and FSCV measurements, the CFME was placed in the FUS prototype through the 2 mm operative channel described above, and it was fixed using a piece of reusable adhesive putty (Fig. 2(e)).

### d. DA release neuromonitoring by FSCV: CFME fabrication

A custom-made CFME was fabricated to be embedded in the FUS neurostimulation platform, to provide FSCV neuromonitoring of FUS-evoked DA releases. Glass capillaries with 1.5 mm outer diameter and 0.86 mm inner diameter (Harvard Apparatus, USA) were pulled using a Flaming/Brown heat puller (P-97, Sutter Instrument, USA). Silver conductive paint (Electrolube SCP03, Leicestershire, UK) was used to attach 30 µm carbon fibers (Goodfellow Corporation, PA, USA) to 4 mm diameter copper wires, ensuring electrical contact between the CFME and the recording device. After the Silver paint dried, copper wires and carbon fibers were inserted into the already prepared glass capillaries. The sharp tips of the capillaries were dipped in Epoxy 301 (Epoxy Technology, Inc., USA) and connected to a vacuum pump to aspirate the epoxy resin before drying at 57 ^◦^C for 24 h. To expose the carbon fiber surface, the CFMEs were polished at 90 ^◦^ angle with a homemade polishing device covered with diamond lapping sheets with a grit size of 3 µm (LF3D, Thorlabs, Inc., USA), generating a 2-dimensional polished disc-shaped carbon surface (Fig. 2(b, c)). Then, Nafion (5 wt% in alcohol/water, Ion Power, Inc., USA), a perfluorinated ion-exchange polymer, was electrodeposited on the carbon fiber surface for 30 seconds by applying a constant voltage of 1V versus an Ag/AgCl reference electrode. Following electrodeposition, the electrodes were left to dry for 24 hours^21,24^ at room temperature.

A current-to-voltage amplifier (VA-10x, NPI Electronic, Germany) and a Multifunction IO device (USB-6341, National Instruments, Austin, TX, USA) were utilized for FSCV recordings. WCCV 4.0 software (Knowmad Technologies LLC, Tucson, AZ, USA) was used to gather and analyze FSCV data. A triangle waveform from −0.4 V to 1.3 V versus Ag/AgCl was scanned at 400 V. s^−1^, which enabled the redox reaction of DA; the procedure involves the oxidation of DA to DA-o-quinone (DAQ), which is then reduced back to DA (Fig. 2(b)). The waveform lasted 8 ms and was applied every 100 ms. Data filtering (zero phase, Butterworth, low-pass, 2 kHz first cutoff) was performed using the WCCV 4.0 software.

FSCV recordings of DA release in cell cultures were performed in DPBS at room temperature.

### e. Simultaneous *Ca*^2+^ activity neuromonitoring by florescence microcopy imaging

*Ca*^2+^ signaling in cell cultures was monitored by fluorescence microscopy imaging simultaneously with monitoring DA release by FSCV. Cells were permeabilized with Pluronic acid (F-127 Pluronic, P6866, Invitrogen, USA), and incubated for 30 min in culture medium containing 3 µM Fluo-4 (ab241082, Abcam, UK) and 3 µM F-127 for standard cells, 0.5 µM Fluo-4 and 0.5 µl.ml^−1^ F-127 for dopaminergic cells. Cells were then incubated 30 min in culture medium (DMEM-F12 for dopaminergic differentiation and complete culture medium in standard cells) before being imaged in DPBS containing calcium. Fluorescence imaging was performed using a 494 nm excitation light (pE-300-W, 1523 CoolLED, UK) and a single-band emission filter (FITC-3540C-000, Semrock, USA). *Ca*^2+^ fluorescence was tracked under ×10 optical magnification by the ORCA-Fusion CMOS Camera (C14440-20UP, Hamamatsu Photonics, Japan); operated at 10 frames per second. *Ca*^2+^ dynamics were recorded for 30 seconds. The application of the FUS pulse was triggered 10 s after the start of acquisition.

### f. Calibration of the CFME-based biosensor

Calibration of the CFMEs was carried out using a 3D printed flow cell previously developed by Hexter et al.^25^. The flow cell was linked to a 6-port HPLC valve (Shimadzu, Japan) and a syringe pump (Vial Pilote A2, Fresenius Vial Infusion Technology, Germany). The flow rate was set at 1.7 ml·min^−1^. The syringe was filled with PBS 10 mM. Sample of DA solution was prepared at various concentrations in PBS 10 mM. The CFME and a pseudo-reference wire of Ag/AgCl were placed in the flow cell and exposed to the PBS solution. The DA FSCV waveform was first applied to the CFME for 10 minutes at 60 Hz, followed by 10 min at 10 Hz. After this initial cycling, a 30 second acquisition began. After 5 s of baseline recording, the HPLC valve was switched to inject the DA solution (DA hydrochloride, Sigma-Aldrich, H8502, Merck, Germany, 158 nM); at t=15 s the HPLC valve was switched back to the PBS flow.

### g. FUS prototype characterization and single-pulse neurostimulation sequences

The impedance spectrum of the FUS transducer was measured using a vector network analyzer (Planar TR1300/1, Copper Mountain Technologies, USA). The fundamental resonant frequency of the transducer was measured to be *f*_0_ = 0.68 MHz. However, the transducer was driven at a higher harmonic frequency in an effort to minimize the size of the focal spot to achieve higher spatial selectivity. Increasing the FUS frequency also allowed promoting *in-vitro* repeatable non-destructive and well-tolerated effects of the mechanical forces directly induced by the FUS field^16^. Using the previous version of FUS implant prototype (no operative channel, no hole in the spherical transducer), and 2 harmonic frequency configurations (*f*_11_ = 8.09 MHz and *f*_7_ = 5.19 MHz) already validated by Aubier et al.^16^ on standard cell culture, preliminary trials were conducted on DA cultures to select a frequency of FUS neurostimlulation.

Since the highest 11^*th*^ harmonic *f*_11_ associated to the finest focal spot (190 µm) was not repeatable or reproducible in inducing responses in DA cells, the 7^*th*^ harmonic *f*_7_ was chosen. For the novel FUS prototype used throughout the present study, this corresponded to *f*_7_ = 5.11 MHz. For pressure field measurements, a certified and pre-calibrated needle hydrophone (0.2 mm, Model 3558, Precision Acoustics Ltd., Dorchester, UK) was used to measure the distribution of acoustic pressure produced by the novel FUS prototype in water (at the steady part of the pulse with a 100% duty cycle). A commercially available computer-driven three-axis motorized hydrophone bench (UMS Test Tank, Precision Acoustics Ltd, Dorchester, UK) was used for 2D mechanical scanning in order to reconstruct pressure maps in longitudinal (x-z and y-z) and transverse (x-y) planes. Since the piezoelectric ceramic is spherical (being symmetric in x-y plane), only x-z plane was plotted. The pressure field measurement was done at low signal levels (low pressures) with the objective of studying the impact of including the tip of the CFME in the middle of the acoustic field created at the 7^*th*^ harmonic and comparing it to the free-field circumstances; considered as a water tank with no Petri dish, cells, PDMS, or other obstacles. These low-pressure measurements focused on the relative fluctuations caused by the CFME rather than the absolute pressure values. The CFME was placed 5-6 mm away from the focus point to reduce the risk of interaction with the hydrophone.

The output acoustic powers emitted by the FUS prototype at both the fundamental (*f*_0_) and 7^*th*^ harmonic (*f*_7_) frequencies were determined as a function of the input driving voltages using the radiation force balance (RFB) measurement method and a commercially available RFB instrument (RFB 2000, Onda Corporation, USA). These measurements were performed ten times and averaged for each increment of driving voltage. To identify the FUS stimulation sequences responsible for the *Ca*^2+^ response, we used Aubier et al.^16^‘s parameters, which consistently activate *Ca*^2+^ responses in control cells. To determine the minimum threshold for triggering a *Ca*^2+^ response in dopaminergic neurons, we started with these parameters and utilized a pressure-escalation technique; FUS stimulation was thus performed on different areas of a neuronal culture while gradually increasing pressure levels. We carried out the escalation directly on dopaminergic neurons and then confirmed that the improved protocols were effective and safe for both cell types, with no apparent cellular harm. This procedure resulted in the determination of the 700 µs single-pulse duration and the minimum spatial peak RMS pressure (*p*_*spr*_), which represents the minimum effective stimulus shared by both populations. We chose a single-pulse protocol because, as previously stated in the literature^17,26^, low-energy FUS sequences can modulate neuronal activity without causing irreversible damage, and single, higher-intensity pulses provide a simpler framework for characterizing fundamental neuromodulatory effects before progressing to more complex repetitive-pulse protocols (Fig. 3(a)).

**Figure 3.**
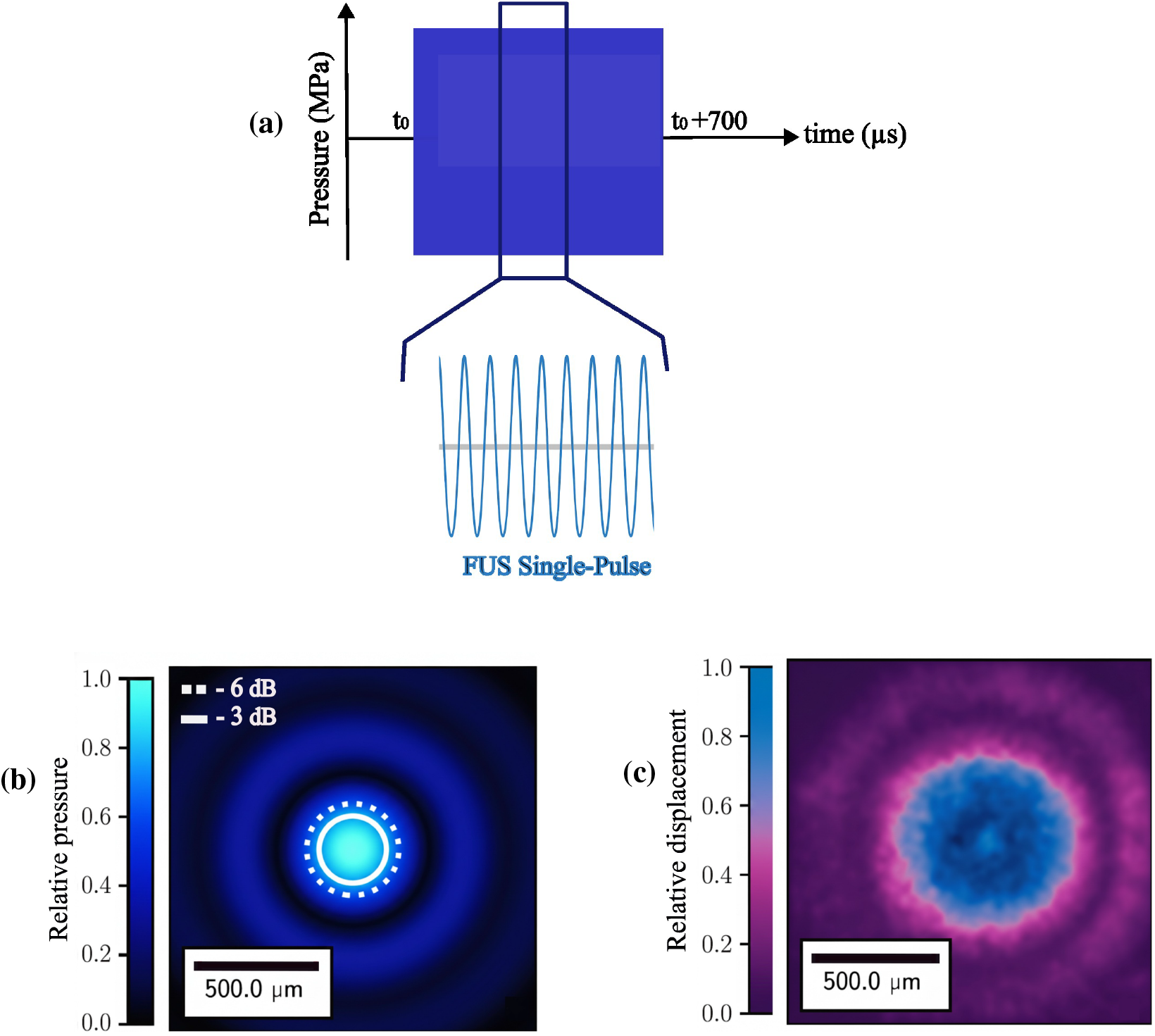
Characterization of single-pulse focused ultrasound (FUS) stimulation. (a) Schematic representation of the 700 µs single-pulse FUS stimulation sequence. (b) Numerically simulated spatial distribution of ultrasound pressure generated by the single-pulse FUS stimulation. (c) Qualitative visualization of FUS-induced mechanical stresses in an optical phantom, assessed using microscopy imaging combined with optical particle tracking methods.

### h. Numerical simulation of FUS-induced pressure field and heating *in-vitro*

Acoustic and thermal simulations of ultrasonic propagation and heat deposition in a multilayer experimental setup were carried out numerically using the k-Wave MATLAB toolbox^27^. The simulation was built on the k-space pseudo spectral approach, which numerically solves first-order acoustic wave equations in both space and time while accounting for frequency-dependent acoustic absorption and boundary constraints. A spherically focused transducer (15 mm aperture, 15 mm radius of curvature) was modelled as a curved source surface with a 2 mm central hole. The radiated acoustic power sufficient to induce a Ca^2+^ response at *f*_7_= 5.11 MHz, measured by RFB, was converted to a pressure boundary condition using the following equation:

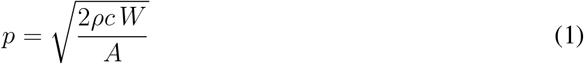

where *p* is the acoustic pressure amplitude, *ρ* and *c* are the density and sound speed of the coupling medium, *W* is the acoustic power, and *A* is the transducer’s effective surface area. The computational domain included multiple layers—air, agarose coupling gel, 3D printed coupling cone, water, glass coverslip, and PDMS—each assigned appropriate density, speed of sound, and attenuation parameters from literature values. Perfectly matched layers were used to minimize reflections at the domain boundaries. The spatial distribution of the acoustic pressure amplitude and its on-axis profile were extracted to identify the focal region (Fig. 3(b)). Subsequently, the Bio-Heat Transfer Equation (BHTE) was solved to evaluate temperature elevation:

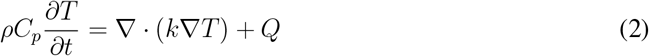

where *ρ* is density, *C*_*p*_ is specific heat, *k* is thermal conductivity, and *Q* = 2*αI* represents the absorbed acoustic energy per unit volume (with *α* being the absorption coefficient and *I* the acoustic intensity). Temperature distributions were simulated for both FUS on-time (700 µs) and cooling phase (1 s), with initial temperature *T*_0_= 37 °C. The resulting temperature maps and temporal profiles were analyzed at the interface between the water and PDMS layers to assess focal heating under exposure. Temperatures were computed at the PDMS-cell media interface, both in attenuated domain of the PDMS itself and in the cell culture medium. Because the cell medium has little acoustic attenuation (similar to water), direct local absorption of FUS energy should result in insignificant heating. However, considerable heating was expected due to the thermal diffusion of PDMS into the cell medium.

### i. FUS targeting of neural cultures

Before stimulating neural cells, acoustic radiation force-driven displacement measurements on an optical tissue phantom were performed to confirm appropriate FUS targeting. The PDMS-coated Petri dish was filled with a 4 ml solution of water and cornstarch (Maïzena, Unilever, UK). Following the settlement of the starch on the PDMS surface, the particles were illuminated laterally (OZM-A4516, Kern & Sohn, Germany) and utilized as an optical contrast agent to qualitatively determine the spatial distribution of FUS-induced mechanical stresses in the microscope field-of-view (FOV) (Fig. 3(c)). Since PDMS is optically transparent, using cornstarch increased the visibility of surface displacement. This method enabled real-time evaluation of FUS targeting based on particle mobility captured by the live camera. The position of the CFME-containing transducer along the z-axis (ultrasound propagation axis) was adjusted by reducing the diameter of the displaced region while maintaining a safe distance between the CFME fragile tip and the PDMS surface, ensuring that the PDMS surface was centered in the FUS focal zone. The location of the carbon fiber tip was visually verified on the microscope FOV, and its distance to the cell layer varied from 200 µm to 1 mm. Eventually the transducer’s final position was recorded, and it was elevated using electrical micromanipulators. The optical phantom was replaced by a Petri dish containing the cell culture, and the transducer was moved to the previously recorded coordinates.

### j. Data Selection and Analysis Strategy

All *Ca*^2+^ imaging data was reviewed to select films with a clearly apparent fluorescence response after FUS stimulation. Further validation of the films was performed using ImageJ free access software (National Institutes of Health, Bethesda, MD) to confirm the presence and timing of *Ca*^2+^ activation; the first frame was then subtracted from the other frames, and the initial subtracted frame represented the baseline 10 seconds before FUS sonication (*t*_0_-10). In our trials, we collected frames at 1 s (*t*_0_+1), 10 s (*t*_0_+10), and 20 s (*t*_0_+20) after sonication. Signal and image processing were performed on all captured films using in-house developed Python code, as detailed by Aubier et al.^16^. The quantitative *Ca*^2+^ dynamics were extracted, resulting in *(i)* fluorescence-intensity-change plots (Δ*F/F*) and *(ii)* spatiotemporal maps that show the signal’s evolution over time. The *Ca*^2+^ activation event for each cell was represented as red crosses. To consider a fluorescence alteration as an activation event, we picked responses that exceeded 10 times the baseline fluorescence fluctuations. For response characterization, we focused on the first 2.5 s following FUS exposure. This window consists of both the immediate focal response (<1 s), as shown in recent work by Aubier et al.^16^, and the very beginning of the delayed propagative response (for 1.5 s). *Ca*^2+^ responses were compared between standard cells and dopaminergic neurons over this time period. Statistical analyses were carried out in Excel using chi-square test; p-values < 0.05 indicate statistically significant results^28^. To connect optical data with electrochemical measurement, FSCV files for selected *Ca*^2+^ films were studied. Voltammograms were obtained 10 seconds after stimulation (*t*_0_ + 10 s) for six independent samples in dopaminergic and standard cultures. The acoustic exposure conditions were reported using standard quantitative descriptors: *I*_*sppa*_ and *I*_*sapa*_ refer to the spatial-peak pulse-average and spatial-average pulse-average acoustic intensities (W·cm^−2^), respectively. *p*_*sp*_ and *p*_*spr*_ denote the spatial-peak and spatial-peak RMS pressures (Pa). *E*_*total*_ indicates the total quantity of acoustic energy emitted by the transducer active surface. Focal-spot areas were determined based on the acoustic intensity distribution’s full width at half maximum (FWHM, −3 dB). The maximum temperature rise (°C) at the cell surface (Δ*T*_Cell Max_) and the PDMS layer (Δ*T*_PDMS Max_) was determined using k-Wave simulations in MATLAB, using the previously reported bioheat transfer equation.

## Results

### a. Perforated FUS transducer characteristics

The impedance spectrum of the new perforated transducer was measured to determine its fundamental (*f*_0_= 0.69 MHz) and harmonic resonant frequencies. The focal spot diameter for the wave traveling in water, at the fundamental frequency was approximately 2 mm, while at the 7^*th*^ harmonic (*f*_7_= 5.11 MHz), it was reduced to about 293 µm, providing a significantly finer focus; while promoting repeatable non-destructive effects of the mechanical forces directly induced by the FUS field, as previously shown by Aubier et al.^16^. Therefore, the 7^*th*^ harmonic frequency was selected for subsequent experiments. In the absence of the CFME, the FUS spatial peak pressure at the focus was measured to *p*_*sp*_ = 0.15 MPa, whereas the presence of the CFME inside the transducer (5-6 mm away from the focal point) induced a 13% drop (*p*_*sp*_ = 0.13 MPa) (Fig. 4). However, we did not observe a major impact in the shape and size of the focal spot when including the CFME as compared to the focal spot of the free-field measurement. As direct measurements of pressure and temperature elevations on the cell culture for *in-vitro* configuration of neurostimulation (presence of the Petri dish, high signal levels: *p*_sp_ of several MPa) were not feasible, acoustic and thermal simulations of ultrasonic propagation and heat deposition in the multilayer setup were performed numerically. The k-Wave simulation provided detailed spatial maps of the acoustic pressure and temperature distributions across the layered medium. The perforated FUS transducer generated a sharply defined focal zone approximately 15 mm from the transducer surface, achieving the **spatial peak pressure** (*p*_*sp*_) in the range of 26.43 ± 3.12 MPa, corresponding to the values able to activate Ca^2+^ signalling in cultured cells. The **spatial-average root-mean-square pressure** (*p*_sar_) within the focal region − 3 dB was 16.88 ± 1.99 MPa, yielding a **spatial-average pulse-average intensity** (*I*_sapa_) of 19.59 ± 4.12 kW.cm^−2^. Thermal analysis indicated temperature elevations of 2.3 ± 0.46 at the cell culture surface (Δ*T*_Cell Max_) and 6.15 ± 1.28 in the PDMS layer (Δ*T*_PDMS Max_). The numerical simulation of the pressure field in the free-field with no barriers was also performed using the identical acoustic power inputs, revealing a *p*_*sp*_ range of 24.08 ± 2.85 MPa in the absence of CFME. All measured and simulated numerical values are detailed in Table 1. Together, these measurements and simulations provided a robust characterization of the acoustic field, ensuring precise control of the ultrasound exposure applied in subsequent biological experiments.

**Table 1:** Acoustic and thermal parameters for the different FUS configurations (in the absence of CFME).

| Parameters | Free-field | *in-vitro* |
| --- | --- | --- |
| $-3$ dB lateral diameter ($\mu\text{m}$) | 293 | 293 |
| $p_{sp}$ (MPa) | $24.08 \pm 2.85$ | $26.43 \pm 3.12$ |
| $p_{spr}$ (MPa) | $17.02 \pm 2.01$ | $18.69 \pm 2.21$ |
| $p_{sar}$ (MPa) | $15.51 \pm 1.83$ | $16.88 \pm 1.99$ |
| $I_{sppa}$ ($\text{kW.cm}^{-2}$) | $19.94 \pm 4.19$ | $24.02 \pm 5.04$ |
| $I_{sapa}$ ($\text{kW.cm}^{-2}$) | $16.54 \pm 3.47$ | $19.59 \pm 4.12$ |
| $E_{\text{total}}$ (mJ) | - | $15.55 \pm 3.26$ |
| $\Delta T_{\text{cell Max}}$ ($^{\circ}\text{C}$) | - | $2.3 \pm 0.46$ |
| $\Delta T_{\text{PDMS Max}}$ ($^{\circ}\text{C}$) | - | $6.15 \pm 1.28$ |

**Figure 4.**
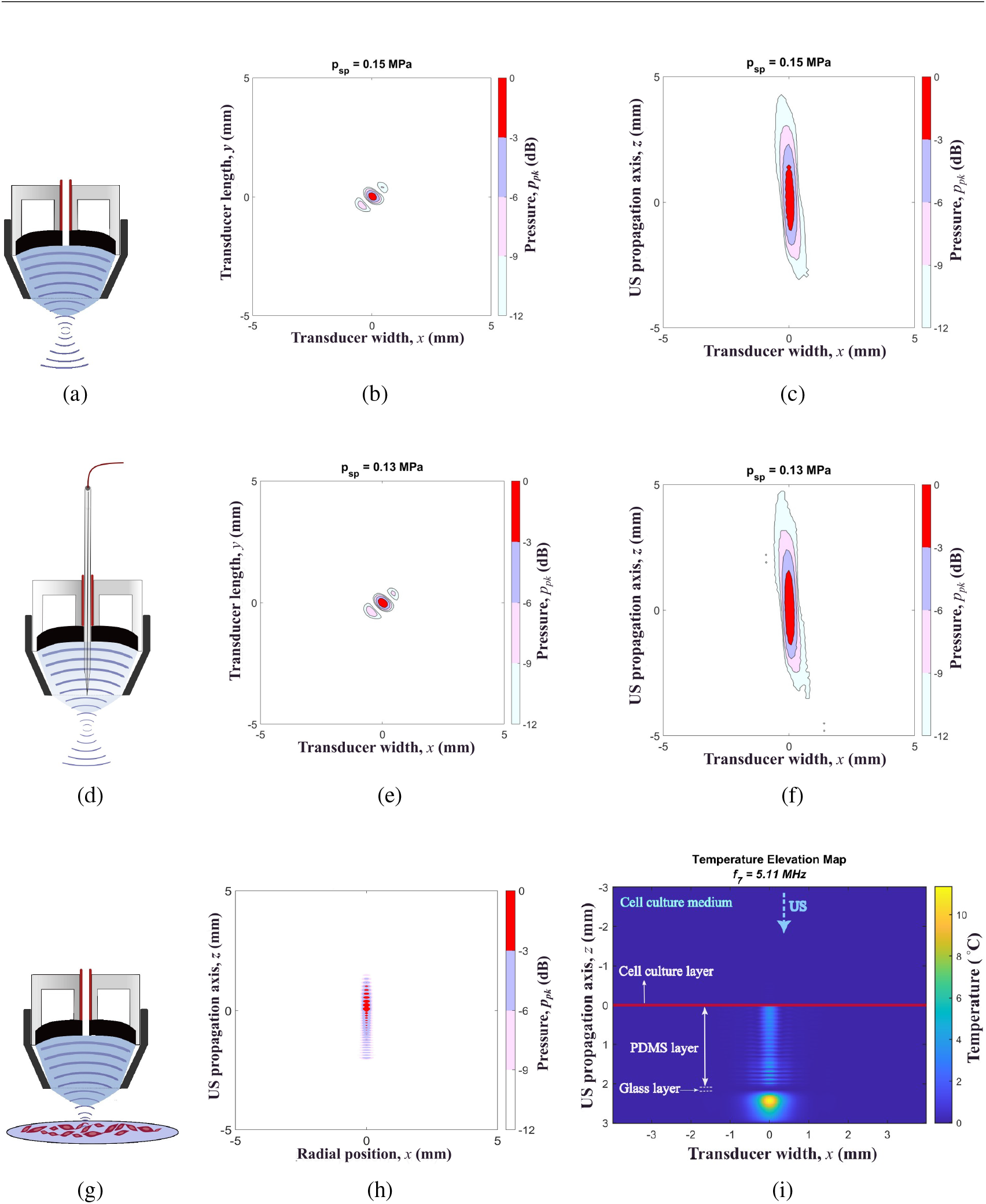
Influence of embedded CFME and petri dish on the FUS field. (a) Cross-sectional view of the transducer without CFME, illustrating FUS propagation in the free field. (b, c) Normalized FUS pressure distribution on the x-y and x-z planes, respectively, with a peak spatial pressure (*p*_*sp*_) of 0.15 MPa in the absence of CFME. (d) Cross-sectional view of the transducer with CFME inserted 5-6 mm away from the focal point, showing FUS emission in the free field. (e, f) Normalized FUS pressure distribution on the x-y and x-z planes, respectively, with CFME present, showing a *p*_*sp*_ of 0.13 MPa. (g) Cross-sectional view of the transducer without CFME, showing FUS propagation within a petri dish. k-Wave simulations were conducted to evaluate the acoustic safety profile for the cells by modeling the pressure distribution and temperature rise within the multilayer experimental setup. (h) Normalized simulated FUS pressure distribution on the x-z plane, with *p*_*sp*_ equal to 26.43 ± 3.12 MPa in the absence of CFME. The simulated acoustic pressure magnitude corresponds to the radiated acoustic power that evoked calcium responses. The plot illustrates pressure variation at the FUS focal point (x, z = 0). (i) Simulated temperature elevation profile within the cell medium, cell layer, and PDMS substrate.

### b. Dopaminergic differentiation of the ReNCell® VM cell line

First, we verified the dopaminergic nature of the differentiated ReNCell® VM cell line. Immunocytochemical analysis was performed using antibodies against tyrosine hydroxylase (TH), the DA-synthesizing enzyme, Nestin, a neuronal intermediate filament protein, and glial fibrillary acidic protein (GFAP), an astrocytic marker^29^.

The ReNCell® VM differentiated cells showed strong TH and Nestin immunoreactivity, whereas GFAP staining was undetectable, confirming that the culture was enriched in dopaminergic neurons (Fig. 5). By contrast, standard differentiated ReNCell® VM (standard cells) showed strong Nestin and GFAP immunoreactivity, but undetectable TH staining. These results indicate that the standard cultures predominantly contained glia and non-dopaminergic neurons, whereas the differentiated cultures were enriched in dopaminergic neurons with minimal glial presence.

**Figure 5.**
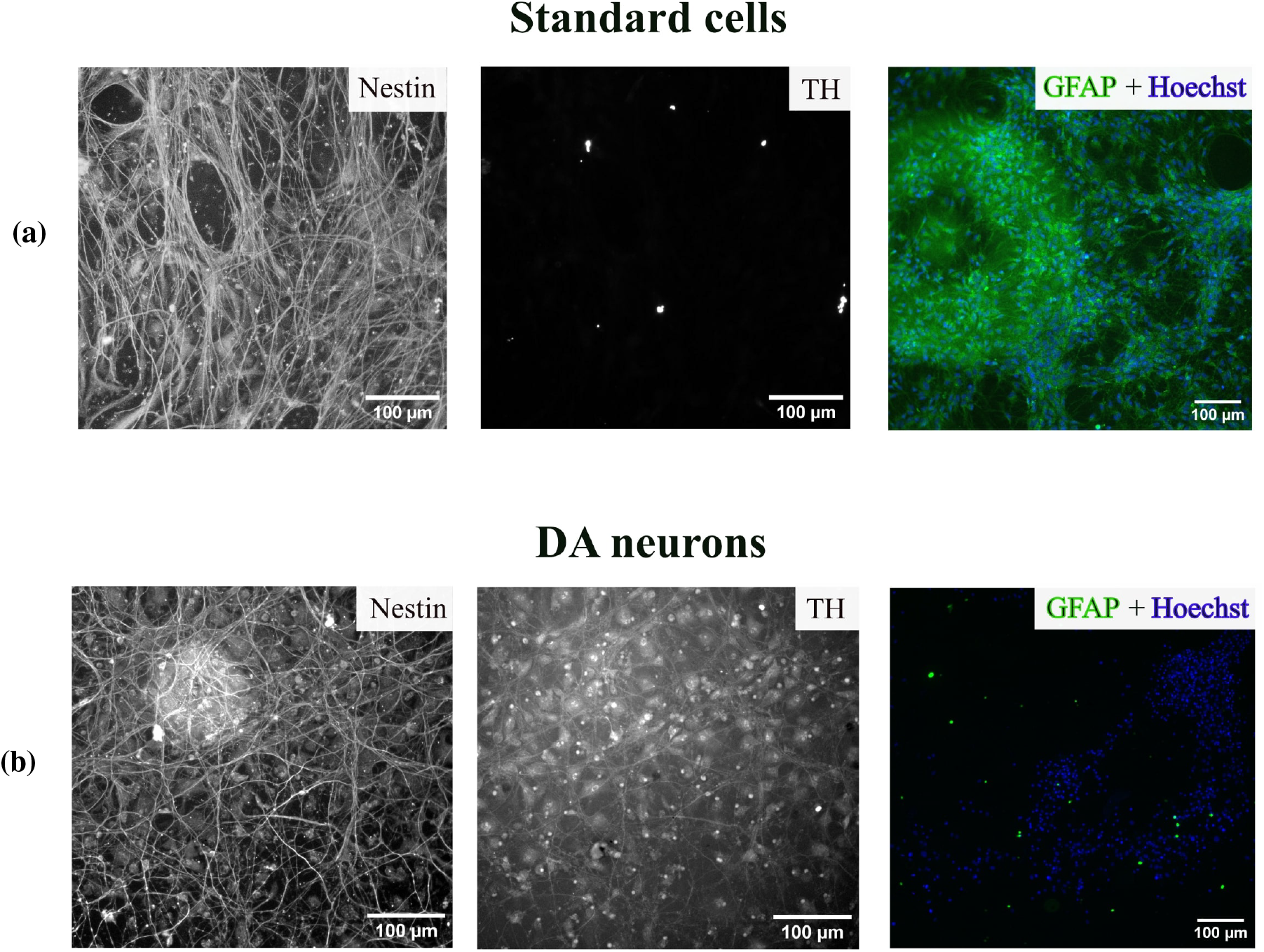
Immunocytochemical characterization of standard cells (a) and dopaminergic (DA) differentiated neurons (b). From left to right in each panel: Nestin, Tyrosine hydroxylase (TH), and GFAP+Hoechst. Nestin: positive staining in both cell types, confirming the presence of neurons. TH: absent in standard cells and present in DA neurons, confirming successful differentiation into DA neurons. GFAP+ Hoechst stainings: present in standard cells and absent in DA neurons, indicating the presence of astrocytes in standard cultures and absence in DA neurons.

The differentiated ReNCell® VM cell line was then stimulated chemically using a high concentration KCl solution to stimulate neurotransmitter release. A CFME was positioned in close proximity to the cell culture, without direct contact with the cell monolayer, and KCl was injected into the Petri dish containing DPBS, reaching the final KCl concentration of 50 mM while FSCV recordings were performed. Cyclic voltammograms acquired immediately after KCl injection revealed a typical dopaminergic electrochemical signature, characterized by an oxidation peak around 0.6 V Vs Ag/AgCl, and a reduction peak around −0.2 V^30^. This signal was also consistent with the results of our CFME calibration utilizing a flow cell following the injection of 158 nM DA into the system.

This dopaminergic signature slowly disappeared 5 and 10 min after KCl stimulation (Fig. 6). The measured current was maximal immediately after KCl administration, reaching about 50 nA at the oxidation peak, and 40 nA at the reduction peak. These results confirmed that the differentiated ReNCell ® VM cell line were indeed enriched in DA neurons with the ability of releasing DA upon chemical stimulation.

**Figure 6.**
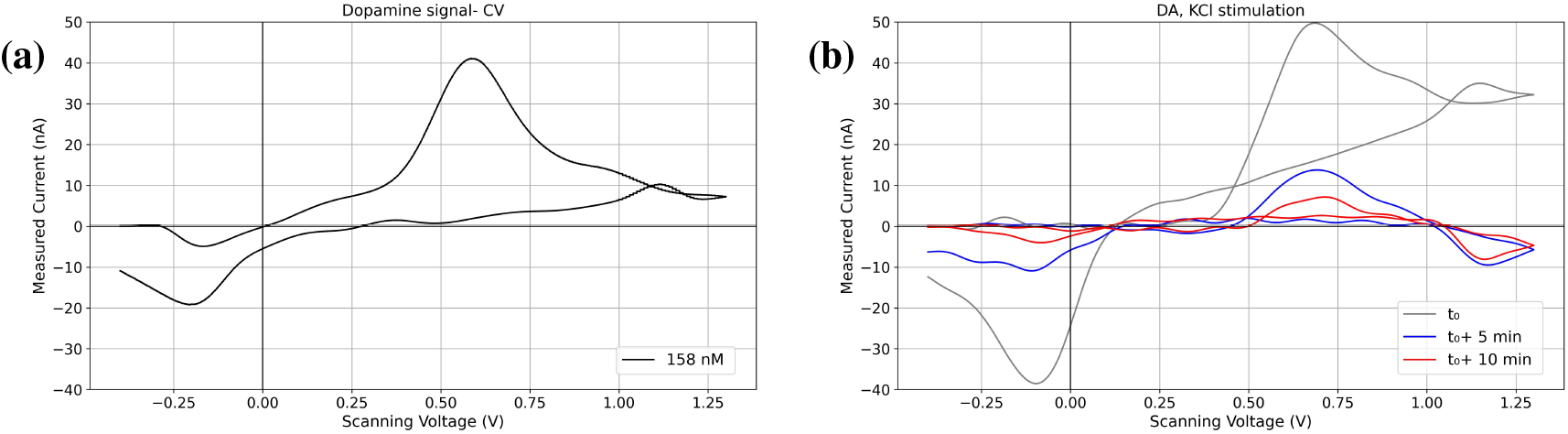
*In-vitro* FSCV DA detection. (a) Cyclic voltammogram (CV) of a DA standard solution (158 nM), showing the characteristic redox peaks of DA. (b) Representative CV plots recorded at stimulation onset (*t*_0_), 5 minutes (*t*_0_ + 5), and 10 minutes (*t*_0_ + 10) after KCl-induced stimulation of DA neurons.

### c. *Ca*^2+^ activities evoked in DA neurons by single-pulse FUS neurostimulation

We then stimulated cultured DA neurons using FUS. The ability of FUS delivered by our novel perforated transducer to elicit *Ca*^2+^ responses was initially evaluated. The higher harmonic frequency of *f*_7_ = 5.11 MHz was chosen to achieve reproducible spatially selective stimulation. Single 700 µs FUS pulses applied at pulse average acoustic powers (*P*_*ac*_) ranging from 12 W to 24 W, were experimentally identified as capable of reliably triggering Ca^2+^ responses. Changes in Ca^2+^ fluorescence intensity of individual cells across the FOV were analyzed over time using images extracted from a 30-s recording, in which the first frame was subtracted from all subsequent frames. In both DA neurons and standard cells, the majority of cells immediately (< 1 s) responding to the stimulation were located within the theoretical − 3 dB surface area (293 µm) of the FUS focal spot (indicated by the yellow circles), consistent with the focal nature of the FUS-induced responses (Fig. 7). Composite representations were created of the spatiotemporal dynamics of Ca^2+^ signaling following FUS stimulation, captured over a 15 s period (showing one representative trial per cell type) (Fig. 7(c, d)). Warm colors correspond to early responses (< 3 s), whereas cooler tones indicate delayed Ca^2+^ activations (> 3 s).

**Figure 7.**
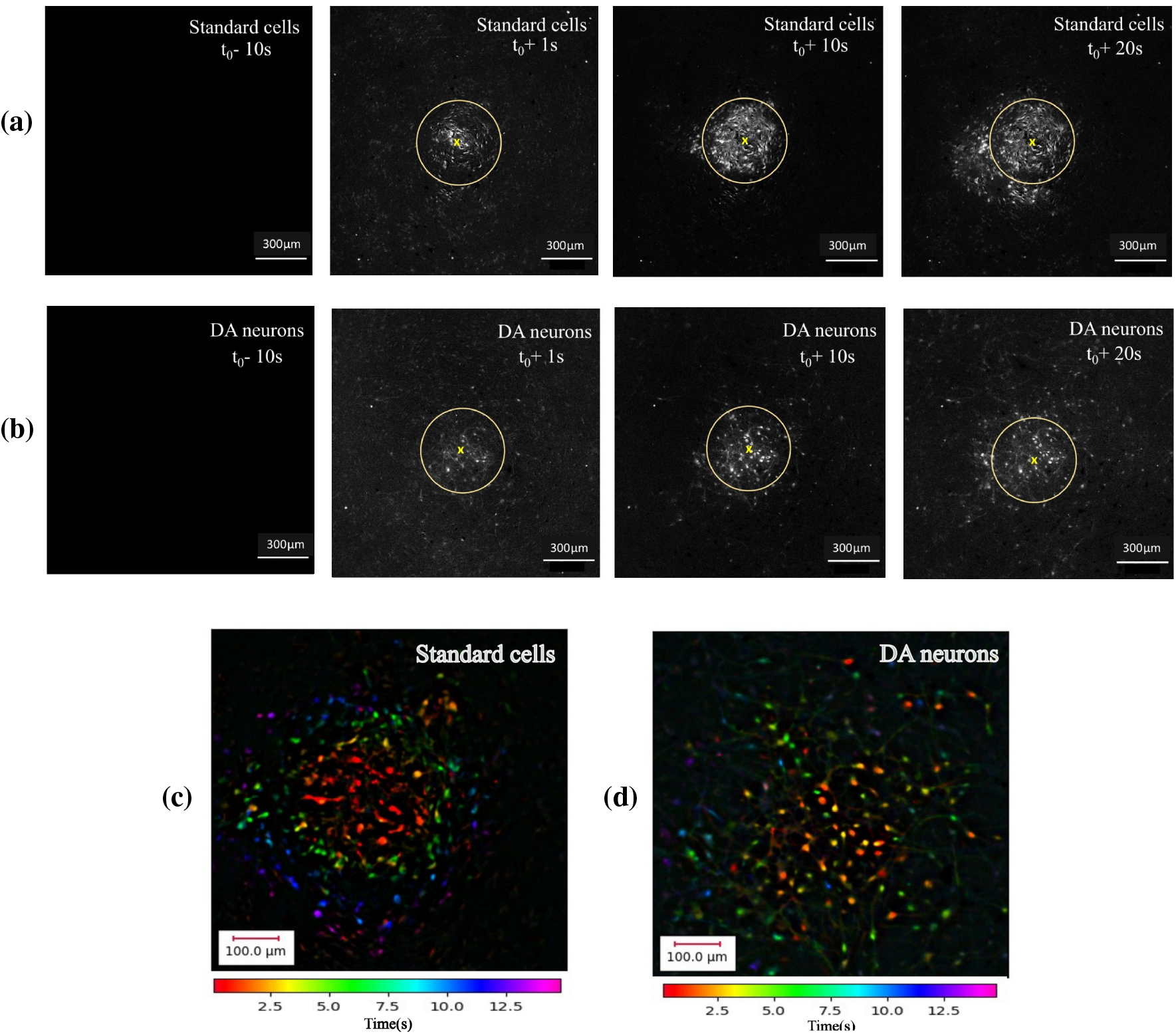
FUS-evoked *Ca*^2+^ responses: spatio-temporal dynamics in standard cells and DA neurons. (a) Representative *Ca*^2+^ response in a standard cell culture at different time points: 10 s before (*t*_0_ − 10 s), 1 s after (*t*_0_+ 1 s), 10 s (*t*_0_+ 10 s), and 20 s (*t*_0_+ 20 s) following FUS stimulation (*t*_0_), showing immediate and omnidirectional propagative activation. (b) Representative *Ca*^2+^ response in DA neurons at the same time points, demonstrating immediate activation with limited and dispersed *Ca*^2+^ responses. (c, d) Spatio-temporal maps of FUS-induced *Ca*^2+^ responses: standard cells exhibit both immediate (< 1 s, warm colors) and propagative (> 1 s, cool colors) *Ca*^2+^ activation, whereas DA neurons show immediate activation with spatially dispersed responses.

In **standard cell cultures**, cells located near the FUS focal point exhibited rapid *Ca*^2+^ activation immediately after sonication (< 1 s), followed by a continuous, ring-shaped omnidirectional propagation extending toward the periphery within the 20 s window (Fig. 7(a)). This pattern reflected a coordinated focal initiation and time-dependent outward spread of *Ca*^2+^ signals. In contrast, **DA neurons** exhibited immediate (< 1 s) *Ca*^2+^ activations not only at the focal area but also at more distant sites from the epicenter. However, the subsequent propagation was limited (no monotonous increase of the activation radius versus time of appearance) and spatially dispersed compared with the standard cell culture (Fig. 7(b)). *Ca*^2+^ activity evoked by single-pulse FUS on standard cells was consistent with that reported by Aubier et al.^16^. However, we observed that the propagation of the *Ca*^2+^ activity triggered in standard cells is substantially wider and more visible than that in DA neurons. FUS stimulation was more effective at the focal region, where the acoustic energy was concentrated. The intracellular *Ca*^2+^ activation responses were analyzed in time and space for both standard cells and DA neurons (Fig. 8). Each activation event was indicated by a red cross, representing the response delay in a given neural cell with respect to the time of FUS stimulation in the x-axis, along with its respective distance from the response epicenter in the y-axis. The delayed *Ca*^2+^ responses exhibited characteristics consistent with a 2-dimensional diffusion process, in which the propagation velocity followed an inverse radial dependence (1/*r*, where *r* is the distance from the epicenter). This spatiotemporal propagation was modeled as a Bernoulli random walk^31^, parameterized by Φ (the diameter of the immediate focal response) and a constant surface expansion rate *e*^16^.

**Figure 8.**
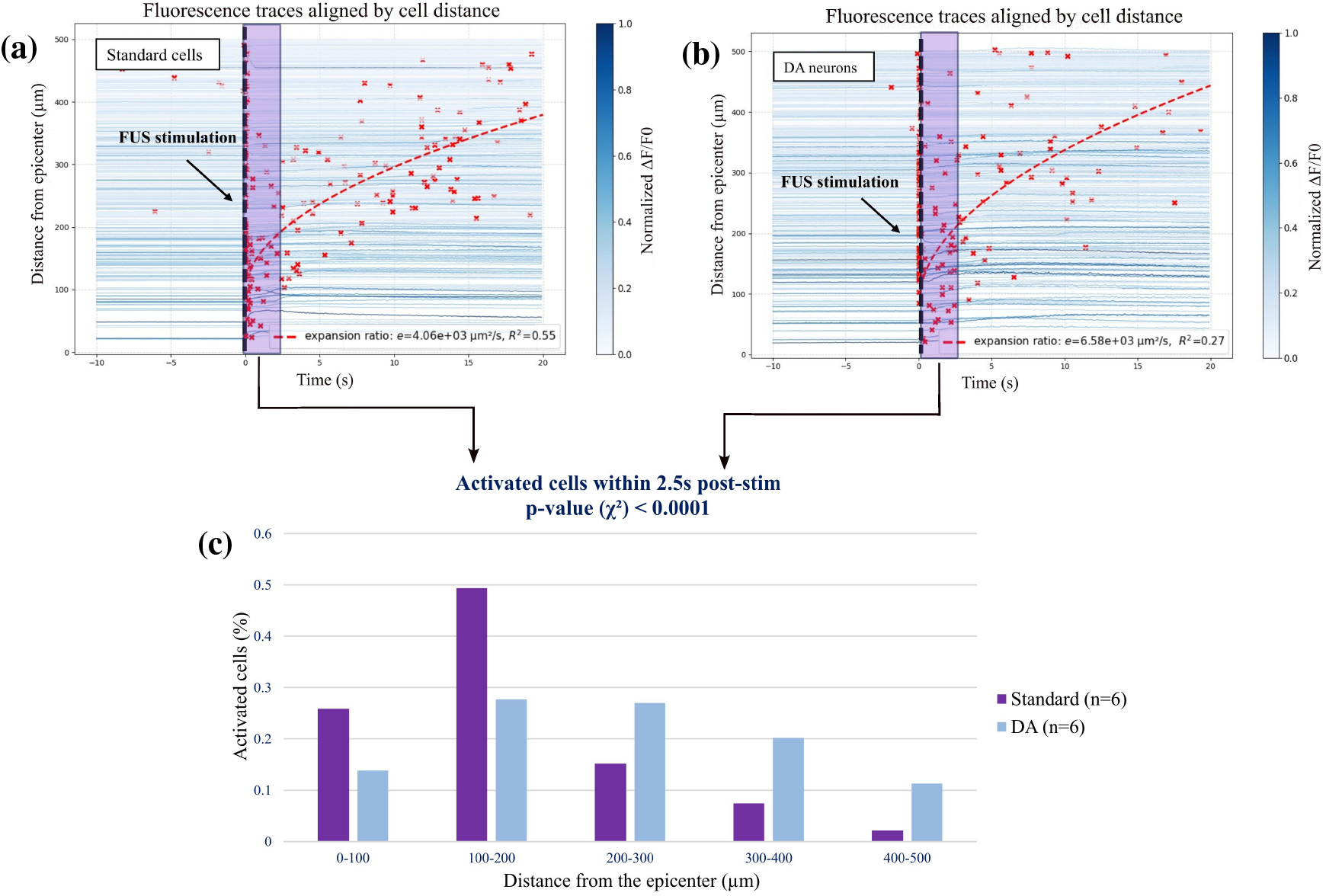
Differences in *Ca*^2+^ spatiotemporal dynamics between standard cells and DA neurons. (a, b) Fluorescence intensity traces (red crosses) showing *Ca*^2+^ activation of cells located at various distances from the FUS epicenter over time. The estimated expansion rates were 4.06 *×* 10^3^ µm^2^. s^−1^ for standard cells and 6.58 *×* 10^3^ µm^2^. s^−1^ for DA neurons; the DA fit had *R*^2^ = 0.27, interpreted as a trend toward faster, more distant activation rather than a definitive linear expansion. (c) Comparison of the number of activated cells in DA neuron and standard cell cultures averaged over six independent trials (n = 6) within a 2.5 s window following FUS stimulation. Standard cells exhibited greater activation near the epicenter (90% for distance< 300 µm), whereas DA neurons showed higher activation at more distant locations (32% for distance> 300 µm). A chi-square test revealed a statistically significant difference in the spatial distribution of *Ca*^2+^ responses (p < 0.0001).

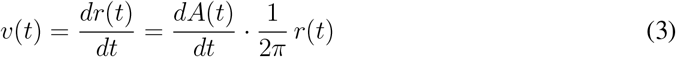

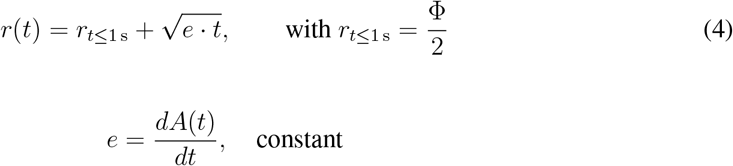

where *r*(*t*) represents the expansion radius as a function of time. *A*(*t*) corresponds to the cumulative response of intracellular *Ca*^2+^ signaling; this response is assumed to arise from a population of neuronal cells responding to neuromodulation over time. Consequently, the rate of change 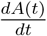 defines the expansion rate of the response region. The fitted model accurately represented the spatial-temporal dynamics of the FUS-evoked responses (Fig. 8(a,b)). Standard cells show a greater coefficient of determination (R^2^= 0.55) compared to DA neurons (R^2^= 0.27), showing that the diffusion-based propagation model in standard cells follows a 2-dimensional diffusion behavior, whereas DA neurons’ responses vary from this.

The mean activation distances within a 2.5 s window following FUS stimulation were averaged across six independent experiments (n= 6) for each cell type, including both the immediate response (< 1 s) and 1.5 s of delayed propagation. Within this period, standard cells exhibited a higher number of activation events within the 300 µm radius of the epicenter, whereas DA neurons showed a greater number of activations at more distant sites. The chi-square test revealed a statistically significant difference between the two distributions with p-value much smaller than 0.05 (Fig. 8(c)). These findings on the cell activation distributions indicate the propagation of *Ca*^2+^ signals within the cell culture was different between dopaminergic and standard cells. While DA neurons displayed a spatially dispersed activation pattern, standard cells showed a regular and concentric propagation model.

### d. DA releases evoked in DA neurons by single-pulse FUS neurostimulation

The ability of single-pulse FUS to evoke DA release from DA neurons was assessed using FSCV with a disc-shaped CFME positioned atop the cell layer and aligned with the center of the FUS beam within the perforated transducer (Fig. 2(e)). The FUS pulse was applied 5 s after starting FSCV acquisition in order to establish a pre-stimulation baseline. Six FUS neurostimulations (n= 6) were performed per comparative groups of DA neurons versus standard cells. Background-subtracted cyclic voltammograms (CVs), representing current versus voltage, were averaged and extracted from FSCV measurements at *t*_0_ + 10 s.

A characteristic DA redox signal was detected in **DA neurons** following FUS stimulation, exhibiting oxidation and reduction peaks (with the currents close to 1 nA and −0.5 nA, respectively), though with smaller amplitudes compared to those evoked by local injection of KCl (Fig. 9(a)).

**Figure 9.**
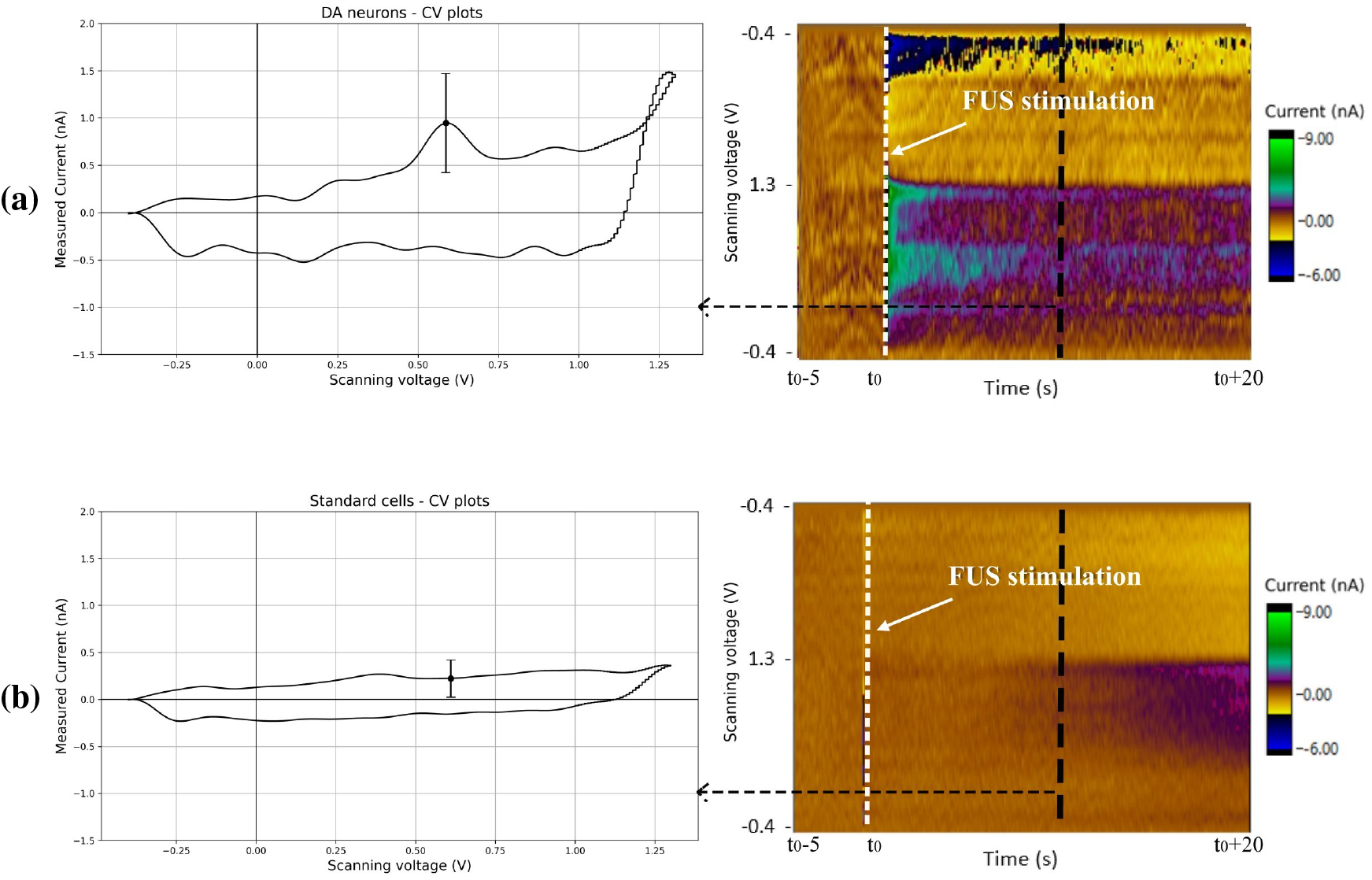
Temporal characterization of FUS-evoked DA release using FSCV. (a) DA neurons. Left: averaged CV recorded 10 s following FUS stimulation of DA neurons (n= 6), showing the characteristic redox peaks of DA. Right: Representative example of FSCV color plot from DA neurons showing temporal changes in oxidation and reduction currents evoked by FUS stimulation. (b) Standard cells. Left: averaged CV recorded after 10 s following FUS stimulation of standard cells (n= 6), showing no DA characteristic peaks, right: Representative FSCV color plot from standard cells showing no detectable current changes over time after FUS stimulation, indicating the absence of FUS-evoked DA release in standard cells.

In contrast, the average CVs obtained from FUS-stimulated **standard cells** did not display the oxidation–reduction peaks characteristic of DA (Fig. 9(b)). The corresponding color plots from representative experiments (Fig. 9(a, b,right)) further illustrate these findings. In DA neurons, the increase of the measured current (green) was observed at the DA oxidation potential, and current decrease (blue) was monitored at the reduction potential, indicating FUS-evoked DA release. Conversely, no such current changes were detected in standard cells, where the background current (yellow) remained stable. These findings indicate that single-pulse FUS stimulation selectively evokes DA release in DA neurons, with no detectable response in standard cells, underscoring the specificity of the stimulation effect.

## Discussion

This study demonstrates that single pulse FUS in the megahertz range (here 5 MHz) can directly activate *Ca*^2+^ signaling from targeted neurons along with neurotransmitter release. To investigate this, we developed a cell culture model capable of differentiating ReNCell progenitors into DA neurons. Immunohistochemical staining confirmed the expression of TH, the DA-synthesizing enzyme, and Nestin, a neuronal intermediate filament protein. The chemical stimulation using a local injection of KCl on these differentiated cultures, led to electrochemically DA release into the culture medium, confirming functional enrichment of DA neurons. In contrast, standard cells were positive for Nestin but negative for TH, indicating the absence of DA differentiation. GFAP staining revealed strong positivity in standard cells but not in DA neurons, confirming that standard cultures were a mixture of neurons and astrocytes. Developing cell cultures predominantly containing DA cells allowed us to directly evaluate the effects of FUS on neuronal populations as opposed to the work done by Aubier et al.^16^ which was done on standard differentiated cells. Indeed, their work identified the observed FUS-induced *Ca*^2+^ propagation dynamic as astrocytic *Ca*^2+^ waves, confirming and complementing observations previously made by Oh et al.^32^.

To monitor DA release evoked by single-pulse megahertz-range FUS stimulation, We designed a novel FUS transducer with a central perforation that allowed insertion of a CFME. This configuration showed minimal interference with the shape of the acoustic field, maintaining the geometry of the focal zone. Comparing to the condition without CFME, the spatial peak pressure (*p*_*sp*_) decreased by approximately 13% when the CFME was present inside the transducer, 5-6 mm away from the focus. This design provides a versatile platform for coupling FUS with various analytical probes such as microelectrodes, biosensors, and microdialysis probes, enabling potential *in-vivo* applications through the CoperniFUS open-source software^33^; which is a flexible Python-based stereotaxic planning platform developed by our team to support FUS treatments through integrated numerical modeling, *in-vivo* targeting workflows, brain atlas registration, and treatment-planning tools.

Using this new design setup, we observed that single-pulse FUS at 5.11 MHz induced rapid *Ca*^2+^ transients in both standard cells and DA neurons. The initial *Ca*^2+^ response was confined to the focal zone but exhibited distinct propagation patterns^16^. In standard cells, propagation was ring-like, while in DA neurons it appeared more dispersed and irregular. DA neurons had wider and dispersed *Ca*^2+^ wave propagation during the first 2.5 s compared to standard cells, which were slower but more coherent. Importantly, in DA neurons —but not in standard cells-FUS-evoked *Ca*^2+^ transients were temporally correlated with electrochemical signals characteristic of DA, showing oxidation and reduction currents at +0.6 V and −0.2 V versus Ag/AgCl, respectively^34^. To our knowledge, no prior *in-vitro* study has directly linked FUS-evoked *Ca*^2+^ transients with DA release from differentiated DA neurons under a single-pulse FUS protocol. Our results bridge this gap by demonstrating, for the first time, that single-pulse FUS can directly stimulate DA neurons, simultaneously eliciting *Ca*^2+^ signaling and electrochemically detected DA release in real time. Previous studies have shown that FUS can modulate neuronal excitability *in-vivo*^35^ and *in-vitro*^36^, modulating calcium dynamics^36,37^, and even behavior *in-vivo* in monkeys^38^ and in humans^39^. Glial cells, which occupy inter-neuronal space and mediate calcium signaling, are known to respond to mechanical stimulation and therefore serve as effective targets for ultrasound-induced activation^40^. FUS has been used to elicit motor and sensory responses in animals and humans^41^, and recent work by Xu et al.^15^ demonstrated that continuous 1 MHz ultrasound can increase DA release—first in PC12 cells *in-vitro*, and subsequently *in-vivo* in rodents. However, unlike this study, which relied on continuous sonication and non-human cell models, our work shows that a single megahertz pulse delivered through a miniaturized implant is sufficient to evoke both *Ca*^2+^ activity and DA release from human DA neurons, quantified in real time using FSCV. This highlights a fundamentally different neuromodulation regime and expands the potential of FUS toward more selective and implant-compatible dopaminergic circuit modulation.

The mechanism by which FUS modulates neuronal activity is complex, involving a delicate interplay of acoustic phenomena at the cellular level. Prior reviews emphasize that acoustic radiation force (ARF)—rather than heating or cavitation—is the primary driver of FUS neuromodulation^42^. The ARF coming from the pressure wave, physically deforms the cell membrane and surrounding structures, likely opening mechanosensitive ion channels, modulating voltage-gated sodium and calcium channels as well as Piezo channels, thereby promoting *Ca*^2+^ influx and neurotransmitter release^43,32,44^. In our study, the application of a single 5 MHz pulse, with energy in the millijoule range, fundamentally shifts the biophysical interaction toward radiation-force–dominated mechanical effects. At this high frequency, cavitation becomes increasingly uncommon, and the single-pulse design prevents cumulative thermal deposition, which is typical in repetitive-pulse approaches. Increasing the FUS frequency also allowed promoting repeatable, non-destructive, and well-tolerated effects of the mechanical forces directly induced by the FUS field *in-vitro*, as shown by Aubier et al.^16^. This frequency-dependent safety profile drastically reduces the probability of destructive stochastic inertial cavitation, enabling the exploration of *in-vitro* FUS pressures beyond current clinical standards. These parameters thus promote high focality, repeatability, and mechanical specificity in stimulation.

Our computational modeling predicts a temporary temperature increase of ∼ 2.3 ± 0.46 °C *in-vitro*, however previous research from our team (Aubier et al.^16^) found no correlation between minimal stimulation thresholds and temperature rise. Furthermore, most temperature-induced effects reported in the literature are modulation or inhibition rather than stimulation^45^. If any thermal influence occurred, it would oppose—rather than facilitate—the responses we elicited. Importantly, this mild heating is mostly a result of the *in-vitro* configuration, including energy absorption by PDMS, and is not expected *in-vivo* settings, where attenuation and perfusion significantly restrict temperature rise. Our approach differs significantly from most previous FUS neuromodulation investigations in that it uses a single-pulse neurostimulation model rather than the more commonly employed repeating low-frequency pulse trains. While our peak pressures are larger, the overall pulse energy remains extremely low (in the mJ range) since the waveform is given once. This varies from shock-wave-based studies, which use significantly higher peak pressures but compress the pulse to a single acoustic cycle^46^; alternatively, our stimulation employs hundreds of high-frequency cycles, allowing for radiation force while using minimal energy. These choices place our work in the rarely explored domain of high-frequency, single-pulse ultrasound neurostimulation.

The temporal coupling we observe between *Ca*^2+^ transients and evoked DA release indicates that FUS triggers stimulated neurotransmitter release, rather than simply altering excitability. Similar *Ca*^2+^-dependent secretory events have been documented in multiple cell types: ultrasound-induced catecholamine release in bovine adrenal chromaffin cells^19^ and *Ca*^2+^-dependent exocytosis in pancreatic *β*-cells^47^. These findings reinforce that *Ca*^2+^ is a central mediator of ultrasound-elicited cellular responses across diverse systems^48^. However, given reports of calcium-independent exocytosis mechanisms^15^ or transporter-mediated DA release^49^, it remains unclear whether DA release observed here was strictly calcium-dependent. Future experiments under calcium-free conditions are needed to clarify this relationship.

The fact that standard cell cultures containing glia displayed ring-like propagation patterns whereas DA neurons showed sparser propagation indicates that the presence or absence of glia in the cell culture can impact the propagation of *Ca*^2+^ signals. This findings are noteworthy for demonstrating that glial cells can engage in FUS-evoked activity and facilitate the propagation in standard cells. The observed responses could thus be the result of both direct neuronal stimulation and glial-mediated network signaling. Interestingly, the acoustic pressures estimated from our simulations are higher than those reported by Aubier et al.^16^. This discrepancy likely reflects methodological differences. The primary objective of this study was to establish a proof of concept for FUS-mediated neural activation and neurotransmitter release, rather than to optimize stimulation thresholds or define minimal safe operating parameters. Accordingly, stimulation conditions were selected to reliably elicit responses, while cell safety was monitored qualitatively by the absence of observable cell detachment and by the ability to repeatedly stimulate the same region. In addition, the incorporation of the CFME within the transducer assembly introduced substantial experimental complexity. Notably, pressure field measurements revealed that the presence of the CFME—although not located at the acoustic focus—can reduce the local pressure by up to 13%, whereas the simulations were performed in its absence. Simulations accounting for the CFME may therefore yield more accurate estimates of the effective pressure range. Despite these uncertainties, comparison of calcium responses in standard mixed cultures and DA neuron cultures suggests that neuronal–glial mixtures represent a more responsive configuration for FUS neuromodulation. Our findings therefore expand the mechanistic landscape by indicating that glial participation is not only possible but may play a fundamental role in shaping network-level activation.

The ability of FUS to consistently trigger *Ca*^2+^-dependent DA release from neuromodulatory neurons has major implications. The DA, serotonin, and norepinephrine systems are key to disorders such as PD and depression. The ability of FUS to evoke neurotransmitter release from neuro-modulatory neurons suggests therapeutic potential for targeting malfunctioning pathways. Indeed, studies using FUS in Parkinsonian models have demonstrated increased DA levels in the medial prefrontal cortex accompanied by behavioral improvements^50^.

While ultrasonic modulation of DA circuits *in-vivo* has been reported^42,51^, the spatiotemporal dynamics of DA release and *Ca*^2+^ signaling at the cellular level have remained poorly understood. By demonstrating single-pulse stimulation–response relationships, our results offer a framework for designing more sophisticated, parameter-tuned FUS neuromodulation protocols. Beyond symptom modulation, controlled DA release may also provide neuroprotective benefits. Excess cytosolic DA can react with oxygen to form reactive species, promoting *α*-synuclein aggregation and dopaminergic cell death in PD. Mosharov et al.^51^ showed that enhanced vesicular trafficking and cytosolic DA clearance protect neurons from oxidative and *α*-synuclein-related toxicity. Thus, our findings support FUS-based neuromodulation therapies, suggesting that FUS-evoked exocytosis and *Ca*^2+^-dependent signaling may not only modulate neurotransmission but also contribute to neuroprotection and neuronal survival in PD and related conditions. However, we should note that *in-vitro* parameters cannot be directly extrapolated to *in-vivo* settings, where vascular, glial, and synaptic interactions play critical roles. Nevertheless, this study establishes a foundational platform for high spatiotemporal resolution FUS neurostimulation/neuromodulation, enabling mechanistic exploration and laying the groundwork for translational therapies targeting DA neurons.

## Conclusion

In sum, we demonstrated that single-pulse, megahertz-range FUS can selectively stimulate human DA neurons, producing rapid *Ca*^2+^ transients and causal DA release in real time. To investigate the underlying mechanisms, we developed a precise *in-vitro* platform that integrates a CFME within a perforated FUS transducer, and we used differentiated DA neuron cultures and control cultures containing both neurons and glia as cellular models.

Our results indicate that FUS-mediated neuromodulation primarily relies on radiation-force–driven mechanical effects, with minimal contributions from thermal or cavitation phenomena. Glial cells were found to participate in propagating network-level responses, highlighting the importance of cellular context in FUS neuromodulation. The single-pulse, high-frequency protocol offers high focality, repeatability, and energy efficiency, distinguishing it from traditional repetitive low-frequency or shock-wave approaches. Moreover, the ability of FUS to trigger *Ca*^2+^-dependent DA release suggests potential not only for targeted neurotransmission stimulation but also for neuroprotective interventions. The compact design of our FUS platform further supports potential transdural implantability, paving the way for customized, minimally invasive therapies targeting disorders such as PD. Overall, these findings provide both mechanistic insights and a translational foundation for precise, single-pulse FUS as a tool for modulating neuromodulatory circuits.

## Acknowledgments

This project was supported by the French National Research Agency (ANR-16-TERC0017, ANR-21-CE19-0007 & ANR-21-CE19-0030), the American Focused Ultrasound Foundation (LabTAU, FUSF Center of Excellence). Additionally, this work was performed within the framework of the LABEX DEV WECAN (ANR-10-LABX-0061) and CORTEX (ANR-11-LABX-0042) of Université de Lyon, within the program “Investissements d’Avenir” (ANR-11-IDEX-0007) operated by the French National Research Agency (ANR). Special thanks to Graduate Initiative NEURO of the University of Lyon I (ANR-21-SFRI0001) who enabled our collaboration with Imperial College London. The authors would also like to acknowledge the contributions and support of the teams involved in this work at the LabTAU (Laboratory of Therapeutic Application of Ultrasound), the CRNL (Centre de Recherche en Neurosciences de Lyon) and the Hashemi lab at Imperial College London.

## Notes

### Competing Interest Statement

The authors have declared no competing interest.

